# Visual Influence Networks in Walking Crowds

**DOI:** 10.1101/2025.01.29.635594

**Authors:** Kei Yoshida, Mario di Bernardo, William H. Warren

## Abstract

Collective motion in human crowds has been understood as a self-organizing phenomenon that is generated from local visual interactions between neighboring pedestrians. To analyze these interactions, we introduce an approach that estimates local influences in observational data on moving human crowds and represents them as spatially-embedded dynamic networks (*visual influence networks*). We analyzed data from a human “swarm” experiment (*N* = 10, 16, 20) in which participants were instructed to walk about the tracking area while staying together as a group. We reconstructed the network every 0.5 seconds using Time-Dependent Delayed Correlation (TDDC). Using novel network measures of local and global leadership (*direct influence* and *branching influence*), we find that both measures strongly depend on an individual’s spatial position within the group, yielding similar but distinctive leadership gradients from the front to the back. There was also a strong linear relationship between individual influence and front-back position in the crowd. The results reveal that influence is concentrated in specific positions in a crowd, a fact that could be exploited by individuals seeking to lead collective crowd motion.

---

Crowds of pedestrians exhibit patterns of coordinated motion in diverse settings, ranging from a narrow platform at a train station to a giant scramble crossing (*1–5*). Similar patterns of movement can be observed in many species, including schools of fish, flocks of birds, and herds of sheep (*6, 7*). Such collective motion has been explained as a self-organizing phenomenon that emerges from local interactions between neighboring individuals (*8, 9*). While many previous studies have adopted a model-based approach that relies on assumptions about individual behaviors and interactions (*10–13*), there is a growing recognition of the importance of a more empirical, data-driven approach (*14, 15*). There are, however, difficulties with trying to infer local interaction rules from global patterns in observational data (*8, 16*). This has led us and other researchers to derive models from experimental data on local interactions (*9, 17, 18*). Here we propose a complementary approach that estimates local influences between pedestrians in observational data and represents crowds as dynamic networks whose nodes correspond to individuals and edges to the visual links between them.

Numerous rule-based models of collective motion have been proposed that define individual interactions based on attraction and repulsion (*11*), heading alignment (*12, 13, 19*), or their combinations (*6, 10, 20*). Most models assume omniscient knowledge about the positions and physical velocities of all neighbors, which is not readily available to individuals embedded in a crowd. To address this limitation, vision-based models have been developed for both humans and animals that simulate group behavior based on the influence exerted on each individual by neighbors within their visual field (*21–26*). The results suggest that crowds and animal groups spontaneously self-organize via visual interactions, exhibiting coordinated pattern formation without external forces or instructions.

We develop a data-driven approach that analyzes pairwise influences in observational data on walking groups. The results are represented as a locally connected graph embedded in a metric space, in which individuals are treated as nodes and the visual influences between them as directed weighted edges. The networks evolve dynamically in time as individuals move about the space. This framework allows us to characterize the collective dynamics by examining both the interactions between individual pedestrians (local level) and the behavior of the entire crowd as a whole (global level) (*27–29*), and ultimately to compare the pattern of observed influences with model predictions.

Previous studies have applied network reconstruction methods to investigate different aspects of organization in animal groups, such as hierarchical structure in pigeon flocks (*30*), directional initiation in migrating white storks (*31*), or communication networks in fish schools (*24*), and to compare metric, topological, Voronoi, and visual networks in a school of golden shiners (*25*). In the context of human crowds, we recently investigated leadership dynamics in behavioral data from small groups of four walking pedestrians (*32*), using Time-Dependent Delayed Correlation (TDDC). Originally developed to determine coordination in pairs of flying bats (*33*), this method compares the heading of one pedestrian with another at all time points and time delays within a temporal window. On each trial, the participants in (*32*) made two turns at unspecified times as they walked together across a room. The TDDC was computed for every pair of pedestrians in the collected position data, which allowed us to infer leader-follower relationships within the group and reconstruct interaction networks at different times during a trial. The results suggested that both spatial position in the group and individual identity play roles in leadership emergence.

However, the interaction networks driving collective motion in human crowds – with specifically visual influences – remain largely unexplored. The present study aims to address this gap by establishing a robust method for network reconstruction and characterizing the structure of influence networks in collectively moving crowds.

We develop an empirical approach that treats human crowds as spatially embedded, visually connected, dynamic graphs, which we term *visual influence networks*. We use pairwise TDDC and the network reconstruction method described in (*32, 34*), which we adapt here by modifying the method of pruning edges to reflect visual occlusion. The resulting networks represent pairwise leader-follower relationships constrained by visibility and visual-locomotor delays. This method is novel in its empirical basis, drawing upon real data from large human groups, and in its consideration of the spatial locations of individual pedestrians within the crowd as well as their visual connectivity. Finally, we introduce novel network measures that estimate the local and global influence of each pedestrian on the rest of the crowd. These measures allow us to characterize the contributions of spatial location, network topology, and individual identity to leadership role in a moving crowd.

Using these methods, we first reconstructed networks in motion-capture data from a human “swarm” experiment and estimated the visual connections. We then investigated the determining factors for leadership emergence in crowds, including the effects of spatial position, network connectivity, and individual identity. We found empirical evidence suggesting that leadership strongly depends on an individual node’s spatial position and its location in the topology of the network.

## Results

The data we analyzed were collected as part of a larger study (*18*). Three groups of participants (*N* = 10, 16, 20) were tested in separate sessions. At the beginning of each trial, participants were positioned at random locations in a starting box. They were instructed to walk about the 14 x 20 m tracking area at a normal speed for 2 minutes, veering left and right while staying together as a group. Motion-capture cameras recorded the head position of each participant, from which time series of heading (travel direction) were computed. This “human swarm” experiment yielded 49 segments of analyzable data with varying durations, a total of 15 minutes of data across the three groups (see sample data in Fig. 1; see Methods for experiment details).

**Fig. 1.**
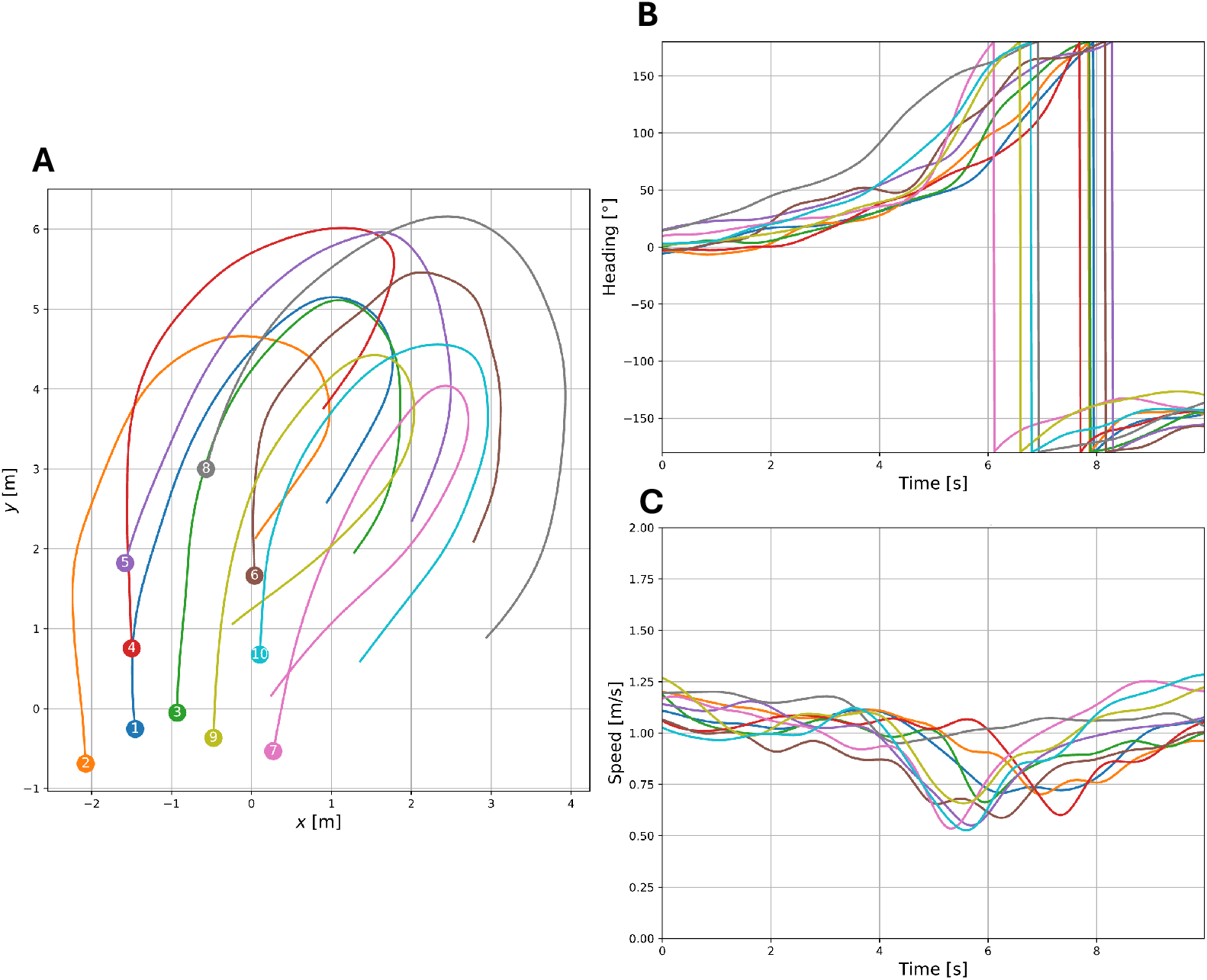
Visualization of a 10-second segment with 10 participants. **(A)** Trajectories of each participant. The *x*- and *y*-axis represent positions in physical space. The starting positions are indicated by the circles containing participant IDs. **(B)** Time series of heading. 0º indicates the positive *y*-axis in panel (a). Positive angles increase clockwise, and negative angles increase counterclockwise from 0º. **(C)** Time series of speed.

### Reconstruction of visual influence networks

To identify leader-follower relationships among pedestrians and reconstruct crowd networks, we computed the Time-Dependent Delayed Directional Correlation (TDDC) from the heading time series for every pair of participants. This analysis compares two trajectories at each time point with various time delays, and assesses the strength of the correlation and direction of influence based on the difference in the heading directions of the two participants. The result is visualized as a TDDC heat map in Fig. 2 (see Methods for a detailed description of TDDC). Pedestrian *i* is considered to lead Pedestrian *j* at time *t* if the time delay *τ* that maximizes the heading correlation is positive (i.e., if the correlation is maximized when comparing the heading of Pedestrian *i* at time *t* with the heading of Pedestrian *j* at time *t+τ*). This is indicated in Fig. 2 by the solid white line (*τ* with the maximum TDDC) being above the black dashed line (*τ* = 0).

**Fig. 2.**
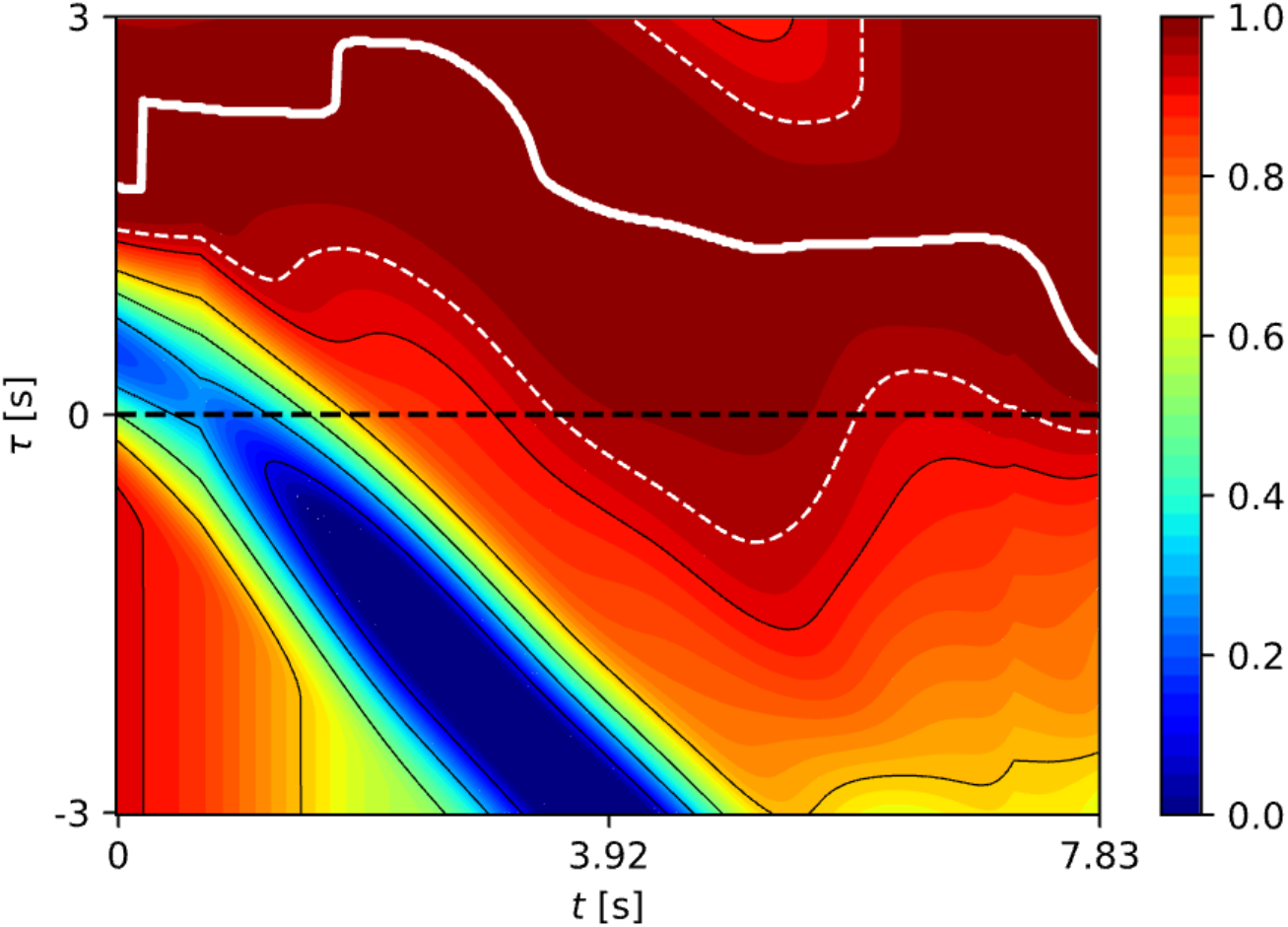
An example heat map characterizing the TDDC values, evaluated for a given pair of participants for a duration of 10 s. The *x*-axis represents time *t*, and the *y*-axis represents time delay *τ*. The color temperature of the heat map represents the TDDC values, where red indicates a value of 1 (i.e., parallel alignment), and blue indicates a TDDC of 0 or below (i.e., there is no meaningful correlation between the two headings). The dashed white contour represents a TDDC value of 0.95, and the dashed black line is where there is no time delay (*τ* = 0). Most importantly, the white solid line represents the *τ*_*ij*_(*t*) value that maximizes the TDDC value as a function of time. At each time *t* where the white solid line is above the black dashed line, it suggests that agent *i* was leading agent *j* at that time. A TDDC map is computed for all possible pairs of pedestrians in a sample of crowd data.

Directed weighted networks were reconstructed for consecutive 0.5-second windows of each segment of data, each visualized as a static “snapshot” of the dynamic network (Fig. 3). In each network, a node represents a pedestrian and a directed edge from node *i* to node *j* indicates that Pedestrian *i* leads pedestrian *j* during the time window. Edge weights, represented by the widths of the arrows, correspond to the percentage of time *i* leads *j* during the window, which we will call the duration of influence. Across the 15 min of analyzed data, 45.84% of all possible directed edges had non-zero weights (see Fig. S1 for the distribution of weights). This suggests that the TDDC captured the directionality of leader-follower relations. An example of the resulting networks over the course of 2.5 s is shown in Fig. 3A (top row), corresponding to the turn in Fig. 1.

**Fig. 3.**
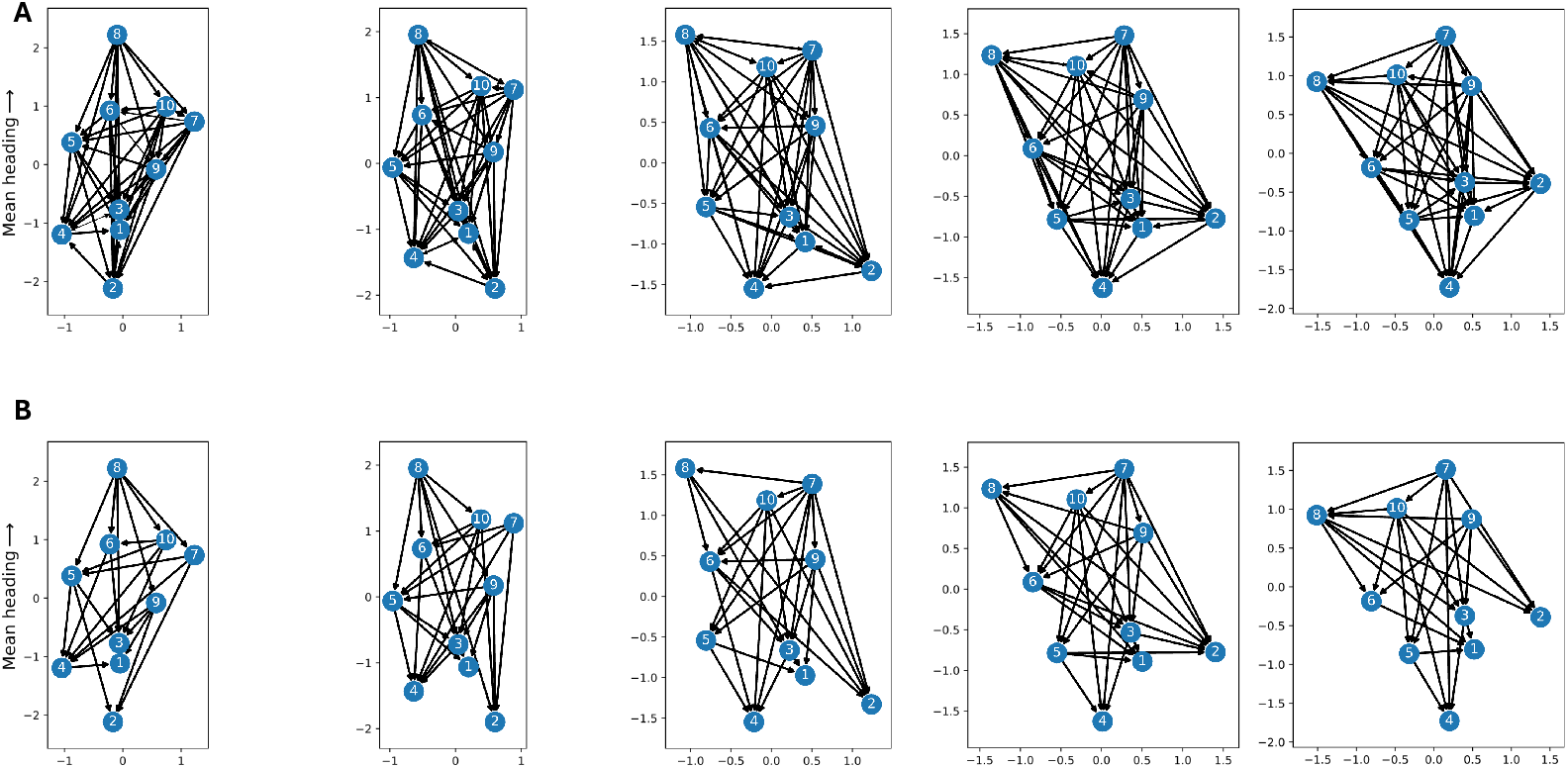
Example network snapshots during a 2.5 s period (*N* = 10). **(A)** Before and **(B)** after pruning methods were applied. Each network was computed for a 0.5 s window, and the series represents a 2.5 s sample from the 10 s segment in Fig. 1 and 4 (from 4.5 - 7.0 s). Participant IDs appear on each node, and node position represents the location of the participant in the crowd, with the group mean heading aligned with the *y*-axis. Arrows on edges represent the direction of influence, and edge weights are indicated by line thickness.

To create *visual influence networks*, edges in each reconstructed network were then pruned based on two criteria. First, the visibility criterion captured the effect of visual occlusion in a crowd: an edge directed from node *i* to node *j* was deleted if, during the 0.5 s time window, the mean proportion of the body width of pedestrian *i* that was visible to pedestrian *j* was less than 0.15 (see Fig. S2). Second, a directed edge was deleted if the mean time delay (with the maximum correlation) was less than a visual-locomotor response time of 300 ms (*35*). As a result of pruning, 51.1% of the initial edges were removed, leaving 22.41% of all possible edges in place. An example of the resulting visual influence networks is shown in the bottom row of Fig. 3, corresponding to the same time intervals as in the top row.

Visual influence networks are directed, weighted, spatially embedded, dynamic graphs. In Fig. 3, the orientation of each network is normalized so that the vertical axis corresponds to the group’s mean heading in the first frame of the time window, and the location of each node corresponds to the spatial position of a pedestrian in that frame. This representation reveals how influence depends on the pedestrian’s spatial position within the crowd and how the network topology evolves in time.

### Network analysis

#### Measures of influence

In each half-second snapshot of the dynamic network, we evaluated the leadership roles of pedestrians using two new measures of influence based on weighted edges. *Direct influence (DI*) estimates the magnitude of a given pedestrian *i*’s influence within their local neighborhood by computing the difference between outgoing edge weights and incoming edge weights for directly connected neighbors. It is defined as:

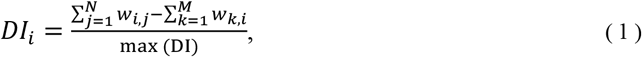

where *j ≠ i* represents the set of *N* pedestrians who are directly influenced by *i*, and *k ≠ i* represents a set of *M* pedestrians who directly influence *i*. The measure takes the sum of the outgoing edge weights *w*_*i*,*j*_ from node *i*, and subtracts the sum of ingoing edge weights *w*_*j*,*i*_ from adjacent nodes, normalized by the maximum DI in the crowd. Thus only incident edges (directly connected to *i*) are taken into account.

*Branching influence (BI*) assesses the global influence of a pedestrian on the rest of the crowd along all outgoing paths in the network, including adjacent nodes and their descendants. It is defined as:

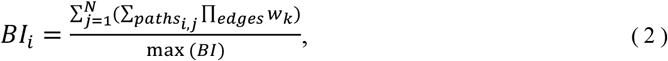

where *j* represents the set of *N* pedestrians who are influenced by *i*, either directly or indirectly, and *k* represents the number of edges along each path linking *i* and *j*. The measure multiplies the edge weights *w*_*k*_ along each outgoing path *i*,*j*, and then takes the sum over the paths. The normalization ensures that the BI value for each pedestrian *i* ranges between 0 and 1. In this case, nodes that are directly connected and indirectly connected to *i* are both taken into account. See Fig. S3 for an illustration.

We also compared these values with two analogous measures of individual influence based solely on degree (number of edges), which do not take edge weights into account. *Direct binary influence (DBI*) evaluates an individual’s local influence by counting the number of outgoing edges from node *i* (outdegree), minus the number of ingoing edges (indegree), and normalizing:

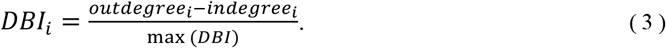

*Branching binary influence (BBI*) evaluates an individual’s global influence by counting the total number of simple paths (with no repeated nodes) originating in node *i*, and normalizing:

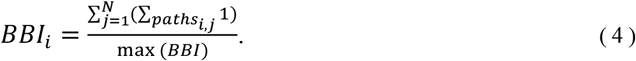

These two measures are equivalent to DI and BI, respectively, with weights set to either 0 or 1. Together, these measures provide a comprehensive understanding of influence and its relation to network connectivity in each snapshot (see Methods for further details).

#### Network dynamics

To investigate network dynamics, we analyzed how leadership changes in successive snapshots. We estimated local and global leadership by ranking the participants according to Direct Influence and Branching Influence in each 0.5 s network (where rank 1 is strongest leadership). Changes in DI and BI during the 10 s of continuous data from Fig. 1 are illustrated in Fig. 4A and 4B, respectively (The corresponding pruned networks appear in Fig. S4). It is clear that rather than being associated with a particular individual, leadership continually shifts as the crowd moves around. Although DI changes on a somewhat faster timescale than BI, they both track the rise and fall of each individual’s influence over the course of the trial. These network dynamics highlight the observation that leadership in a crowd is not static but rather constantly shifts as pedestrians move and their spatial configuration and connectivity change (Fig. S4).

**Fig. 4.**
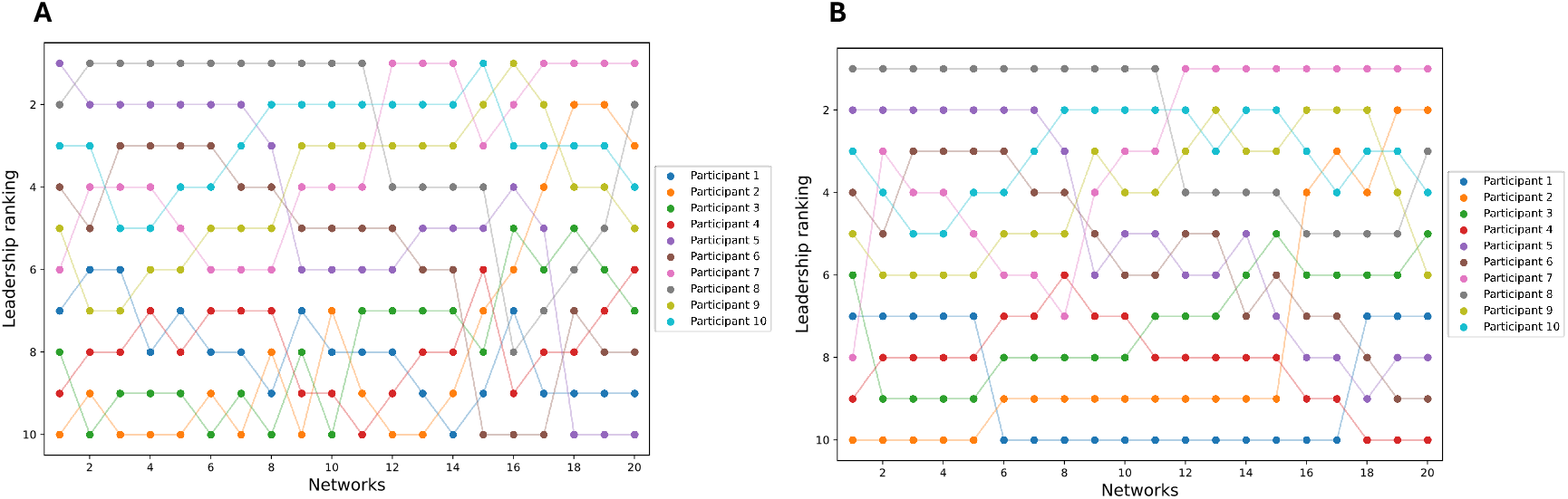
Leadership dynamics of 10 participants showing how leadership ranks change during over 10 s. **(A)** Direct Influence (DI) and **(B)** Branching Influence (BI) ranking. The figure represents the same 10 s segment shown in Fig. 1, and networks 9 to 14 (4.5 s to 7 s) correspond to those in Fig. 3.

To characterize leadership dynamics, we estimated the rate of change in the DI ranking by comparing networks computed in different time windows (0.5 to 4 s) (see Fig. S5 for details). The results indicate that network topology was highly stable over a period of 0.5 s (rank correlation *ρ* = 0.89 between successive 0.5s snapshots), but changed markedly over an interval of 2 s (*ρ* = 0.78 between snapshots 2s apart). Leadership dynamics thus evolve rapidly, but appear to be captured by a 0.5 s snapshot in the present data.

### Leadership depends strongly on spatial position

To investigate how relative position in the crowd affects the strength of leadership, we plotted spatial heat maps illustrating the distribution of influence in the crowd. To account for variations in group size and crowd shape, we first removed spatial outliers (0.91% of all samples) and then normalized each group using the standard deviation (SD) of position on both axes. The vertical axis (Back-Front) was aligned with the group mean heading in each network, and the horizontal axis (Left-Right) was perpendicular (see Supplementary Text); each pedestrian’s position in each snapshot was converted from meters to SDs (a z-score). For each leadership measure, the computed values were averaged across all networks and groups, and then spatial heat maps were plotted with the center of mass of the crowd placed at the origin (0, 0) and the group mean heading aligned with the vertical axis (Fig. 5).

**Fig. 5.**
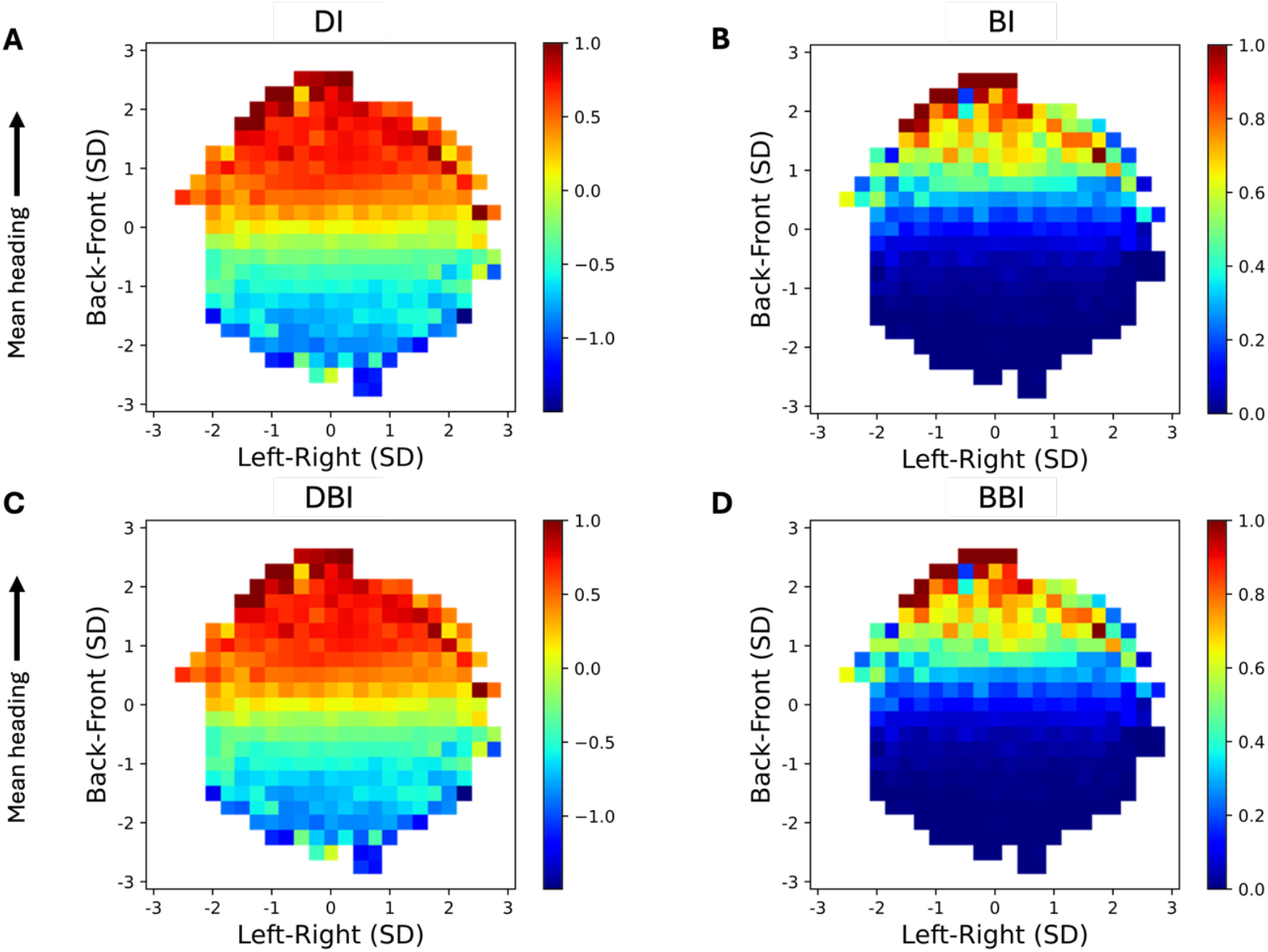
Spatial heat maps of influence, evaluated on the networks after the pruning method was applied. The top row shows measures that take edge weights into account: **(A)** Direct Influence (DI), **(B)** Branching Influence (BI). The bottom row shows measures that are based only on the number of edges: **(C)** Direct Binary Influence (DBI), and **(D)** Branching Binary Influence (BBI). Each cell represents an area of 0.25 SD x 0.25 SD. Color temperature represents the mean leadership value for participants occupying each cell, where red indicates high leadership and blue indicates low leadership.

The normalized spatial heat maps reveal a common pattern of influence across groups. Influence measures based on edge weights (DI and BI) appear in Fig. 5A, 5B, while the corresponding measures based solely on degree (DBI and BBI) appear in Fig. 5C, 5D. Overall, the heat maps reveal a consistent trend in which positions in the front of the crowd tend to have the most influence (warm reds, indicating leaders), while positions in the back of the crowd tend to have the least influence (cool blues, indicating followers). The pattern is the same regardless of group size, as illustrated by separate heatmaps for each group (*N* = 10, 16, 20) prior to spatial normalization (Fig. S6). We also plot the probability density of occupancy in each cell of the normalized heat map in Fig. 6. (For the corresponding occupancy plots for each group, see Fig. S7).

**Fig. 6.**
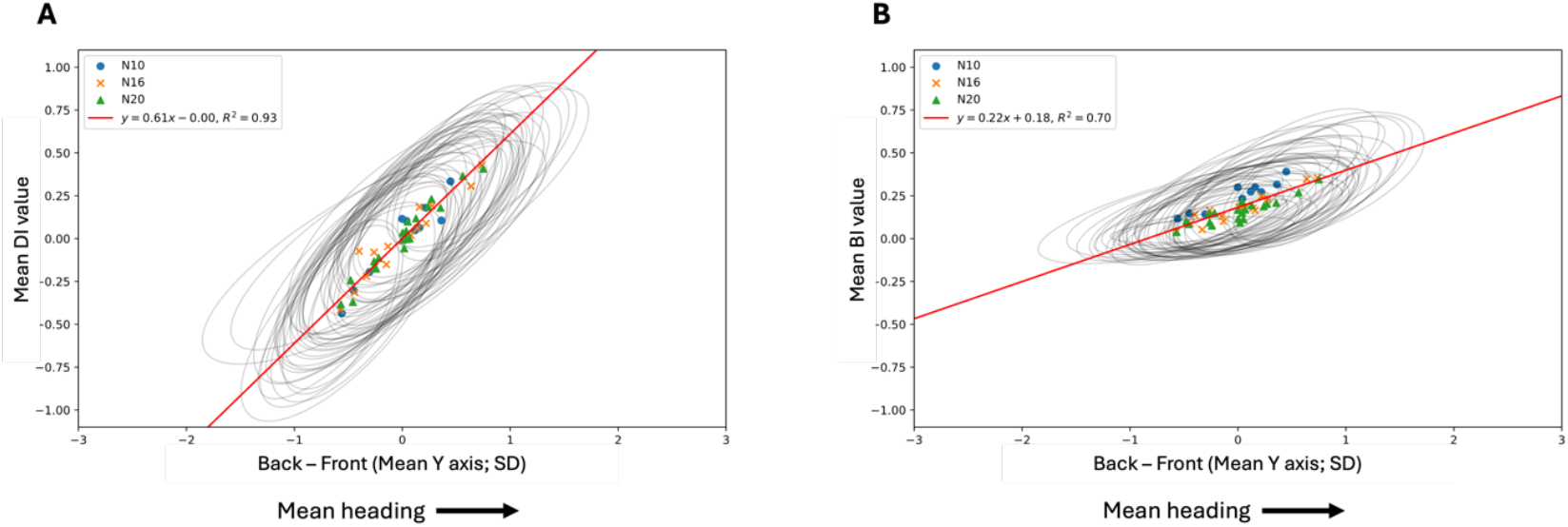
The relationship between individual leadership and spatial position. Individual leadership measured by **(A)** DI (local leadership) and **(B)** BI (global leadership) as a function of the individual’s relative front-back position in the crowd. Each dot represents of the mean of a participant in a given group (blue circles from the *N* = 10 group, orange Xs from the *N* = 16 group, and green triangles from the *N* = 20 group). Error ellipses represent the variation in each participant’s position (radius of 1 SD).

However, the different measures reveal a subtle difference in the gradient of influence from front to back. First, there are distinct differences between local influence (Fig. 5, left column) and global influence (Fig. 5, right column). While direct influence measures (DI and DBI) exhibit a gradual, uniform gradient from front to back, branching influence measures (BI and BBI) reveal a localized hotspot of high influence at the very front of the crowd and a large region of low influence in the rear half. This disparity reflects the different aspects of leadership that are captured by these measures: whereas DI and DBI emphasize local influence within neighborhoods, BI and BBI capture the global influence of each position on the rest of the crowd. Global influence thus flows strongly from the front and is much weaker within the crowd.

Second, there is little difference between influence measures that take edge weights into account (Fig. 5, top row) and those based solely on the number of edges (Fig. 5, bottom row). This result suggests that the observed front-to-back leadership gradient might be attributed solely to the network topology. This suggestion is tempered by two caveats, however. First, network edges depend on neighbor visibility, which is determined by spatial position. Second, edge weights derived from TDDC reflect the duration of influence rather than the strength of influence, which is known to depend on neighbor distance (*18, 21*). Thus, this result does not offer support for a purely topological model of collective motion (*26, 36*). Alternative methods of estimating edge weights will be discussed below.

Third, the effect of visibility pruning is illustrated by comparing the heat maps for DI and DBI after pruning (Fig. 5A and 5C) to those before pruning (see Fig. S8A and S8B). Their similarity indicates that network reconstruction based on TDDC captured the general influence pattern.

Nevertheless, visibility pruning reveals a smoother gradient, with a more homogeneous region of high influence in the front of the crowd shading into a more localized area of low influence at the rear.

### Relationship between individual leadership and spatial position

To explore the relationship between individual leadership and relative position in the crowd, we plotted individual influence as a function of position on the Front-Back axis (which was aligned with the group mean heading) for DI and BI. The results appear in Fig. 6A and 6B, respectively, in which each data point represents a mean influence value and a mean position for one participant in a given group (dot color). Error ellipses with a radius of 1 SD are plotted for each participant.

The analysis revealed a significant correlation between mean influence and mean relative position in the crowd for each participant group (*N* = 10, 16, 20), as well as for all participants combined, for both local DI and global BI measures (all *p* << 0.001). This demonstrates that there is a strong association between an individual’s mean influence and mean relative position in the crowd, regardless of the measure or group. In addition, the major axes of the error ellipses for all participants are oriented along the regression line, indicating that an individual’s influence rose and fell with their front-back position in the crowd.

A regression analysis yielded equations describing the relationship between mean individual leadership (*y*) and mean position (*x*) within the crowd. For DI, the regression equations for the groups with *N* = 10, *N* = 16, and *N* = 20 participants were as follows: *y* = 0.67*x* + 0.00 (*R*^*2*^ = 0.91), *y* = 0.57*x* + 0.00 (*R*^*2*^ = 0.91), and *y* = 0.62*x* – 0.00 (*R*^*2*^ = 0.96), respectively, with an overall regression of *y* = 0.61*x* – 0.00 (*R*^*2*^ = 0.93). This indicates that over 90% of the variance in DI is attributable to spatial position. Similarly, for BI, the regression equations were: *y* = 0.25*x* + 0.25 (*R*^*2*^ = 0.90), *y* = 0.23*x* + 0.17 (*R*^*2*^ = 0.88), and *y* = 0.19*x* + 0.16 (*R*^*2*^ = 0.79) for the respective groups, with an overall regression of *y* = 0.22*x* + 0.18 (*R*^*2*^ = 0.70).

The consistent positive correlations across all groups and leadership measures, combined with the high *R*^*2*^ values obtained in the regression analysis, indicate that a large proportion of the variance in individual influence may be explained by the position occupied by an individual within the crowd. Nonetheless, as indicated by the partially overlapping ellipses in Fig. 6, some individuals reliably occupied positions closer to the front or the rear of the group. This raises the possibility that personal leadership may be expressed by individuals seeking to position themselves near the front of a crowd.

## Discussion

The present study introduces a novel method of reconstructing visual influence networks in moving human crowds, revealing their topological structure and dynamic character. We proposed four network measures that capture local (DI, DBI) and global (BI, BBI) aspects of influence in networks reconstructed from observational data. The analysis enhances our understanding of collective motion by characterizing the visual connectivity between pedestrians and its evolution during crowd movement. Notably, the spatial networks we reconstructed from experimental data demonstrate that influence is concentrated in specific positions within the crowd, primarily due to the network’s topology. To our knowledge, such an investigation has not been previously conducted with human crowds.

The results offer several key insights about collective crowd motion. First, we demonstrate that visual influence networks in moving crowds are dynamic networks that evolve in time, their topology changing on the order of every half-second. Weights, representing the temporal persistence of leader-follower roles between individuals, vary as their positions change, and edges are created and destroyed as neighbors move in and out of view. These network dynamics cannot be predicted from the current topology, however, because they also depend on the individual motions of pedestrians in space. Crowd networks are thus a species of *co-evolving* (or *adaptive*) *networks* (*37*) in which the dynamics of local nodes affects the topology of the network, and the network topology influences the dynamics of the nodes.

Second, the spatial embedding of crowd networks reveals that both local and global influence strongly depends on spatial position in the crowd. The most influential positions are located in the front of the crowd, and the least influential positions are in the rear. There were, however, subtle differences between the influence measures, with local DI exhibiting a gradual gradient from front to back (Fig. 5A) and global BI having a steep gradient from the very front to the center of mass (Fig. 5B). These findings predict that pedestrians will have the strongest influence on crowd movement when positioned near the front of a crowd, a prediction we are currently testing by planting confederates in different locations in a real crowd.

Third, we identified a strong linear relationship between individual leadership and the mean relative position in the crowd for both local and global influence measures. On average across trials, individuals exerting greater influence were identified with positions nearer to the front of the crowd. Although the correlation was high for both local and global influence measures, front-back position predicted local influence more reliably than global influence, accounting for 93% and 70% of the overall variance, respectively. Individual influence also varied with front-back position from trial to trial, as shown by the error ellipses extending along the regression line in Fig. 6. At the same time, different participants tended to occupy positions closer to the front or back of the crowd, a preference that could reflect personal leadership tendencies.

Our method demonstrates the potential of network analysis for understanding collective motion in human crowds, which has only been explored in a few studies of animal groups. Four classes of networks were reconstructed in (*25*) based on different assumptions about interactions in fish schools: metric, topological, Voronoi, and visual networks. By comparing them, the authors found that commonly-assumed metric and topological interactions do not reflect the visual information available to individuals in a school. In another study (*38*), three types of interaction networks were analyzed (metric, topological, and vision network models) based on simulations of fish schools. They were compared using standard network-theory measures and the authors concluded that visual networks are distinctly different from the other two types. We recently reported that visual interactions explain experimental data in humans better than metric or topological interactions (*26*). These studies demonstrate how different assumptions about local-interaction mechanisms generate different network structures and behavioral predictions.

The results of the present spatial analysis (Fig. 5) are particularly informative. Intuitively, given that humans have a 180° field of view, we anticipated that pedestrians at the front of the crowd would influence those at the back, and our results are consistent with that expectation. This finding is not trivial, however – influence in flocks of pigeons, who have a nearly 360° field of view, also flows from front to back (*30*), which may be characteristic of animal groups.

Specifically, we quantitatively identified spatial locations in which local and global leadership may emerge. Whereas local influence (DI) is distributed in a large region in the front half of the crowd and flows smoothly to the back (Fig. 5A), global influence is localized at the very front of the crowd and drops rapidly (Fig. 5B).

While fully disentangling the roles of individual leadership and spatial position in leadership emergence is challenging, we observed a statistically significant relationship between them. This result is consistent with the findings in (*32*) who showed that both positional and personal factors influence leadership. It is difficult to determine the causal role of these factors based on observational data, and future studies should consider an experimental manipulation.

The use of a correlational measure (TDDC) to estimate influence is a potential limitation of our study. The distribution of edge weights is overwhelmingly bimodal with weights of 0 and 1 for various temporal windows (Fig. S1). This suggests that while the TDDC effectively estimates the network connections, the duration of a positive correlation does not adequately quantify the “strength” of influence. Neither does the correlation itself, because it indicates the degree of alignment so its maximum value is usually very close to 1 (Fig. 2). This observation helps to explain why visual pruning only had a small effect on the spatial pattern of influence (compare Fig. 5 left and Fig. S8): pruning eliminated about half the edges, but the effective edges that remained still had strong weights close to 1. It may also explain why the influence measures based on edge weights (DI, BI; Fig. 5, top) differ little from those based on degree alone (DBI, BBI; Fig. 5, bottom). To address this issue, we plan to directly compare TDDC with information-theoretic methods of network reconstruction such as causation entropy (*39–41*).

One implication of the present study is its potential applications to evacuation planning and design. Some scenarios benefit more from clear leadership, as often seen in emergency evacuation (*42*). Our findings suggest that the positions of leaders strongly affect the influence they can have on the movement of the whole crowd. For instance, safety officers may be more effective when they are positioned at the front of a crowd, allowing them to lead pedestrians to safety, as opposed to issuing instructions from the rear. We are currently testing this prediction experimentally.

In conclusion, the present study demonstrates how visual influence networks can be reconstructed from time-series data on real crowd movement. The results highlight how individuals influence the movement of others based on spatial position and visual interactions, consequently driving group-level collective motion.

## Methods

### Human “Swarm” Experiment

The data from this experiment were collected as part of a larger study and have been reported and analyzed in (*18, 26*).

#### Participants

A total of three groups (*N* = 10, *N* = 16, and *N* = 20 participants) were tested in separate sessions. Each group completed four 2-minute trials.

#### Apparatus

The experiments were conducted in Sayles Hall at Brown University. A 14 x 20 m tracking area was marked on the floor with red tape, as well as starting boxes of various sizes. The head position of each participant was tracked using a bicycle helmet with a unique constellation of five reflective markers on 30 – 40 cm stalks. Head positions were recorded with a 16-camera infrared motion capture system (Qualisys Oqus, Buffalo Grove, IL) at a sampling rate of 60 Hz.

#### Procedure

Participants were instructed to walk about the tracking area at a normal speed for 2 minutes, veering randomly left and right while staying together as a group. Participants began each trial in random positions in a starting box to create an initial interpersonal distance (IPD) of approximately 2 m (low-density condition) or 1 m (high-density condition). Each group received four trials, two trials in each of the two density conditions, resulting in a total of 12 trials. They began each trial with a verbal ‘go’ signal and walked around as a group until a ‘stop’ signal.

#### Data processing

A custom algorithm was employed to reconstruct the 3D position of the centroid of the markers on each helmet. The time series of head position in the horizontal (X–Y) plane was filtered using a forward and backward fourth-order low-pass Butterworth filter to reduce occasional tracker error and oscillations due to the step cycle.

The motion capture system failed to track the reflective markers at certain locations in the hall, together with other tracking errors, resulting in some missing data. In this study, we analyzed 49 continuous segments of data with varying durations (1.62 s to 86.97 s) in which the locations of at least 90% of the participants were recovered and at least two networks could be reconstructed based on the selected time window (0.5 s). This resulted in a total of ∼15 min of analyzable data. The IPD conditions were combined for the purpose of this study.

### Time-dependent delayed directional correlation (TDDC)

Time-dependent delayed directional correlation (TDDC) (*33*) measures the leader-follower relationship in pairs of interacting agents by quantifying the alignment of their headings. It compares the heading of one agent with the heading of the other at all time-delays within a certain temporal window. This is based on the assumption that if agent *j* is following another agent *i*, their headings should closely match at some time delay *τ*. In other words, if *i* is influencing *j, j* would adjust its heading to approximate that of *i*.

Given the velocity vector *v*_*k*_ E *R*^*2*^ for agent *k*, the normalized scalar product of the velocity of *i* at time *t* with *j* at time *t + τ* is computed:

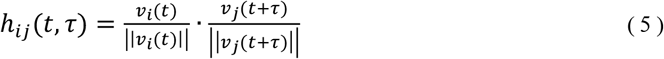

This indicates the degree of alignment between the unit vectors of the two agents at two different times. If *h*_*ij*_(*t*,*τ*) = 1, it indicates parallel alignment of their headings (0°); if *h*_*ij*_(*t*,*τ*) = 0, it indicates an orthogonal relationship in heading directions (90°); and if *h*_*ij*_(*t*,*τ*) = –1, it indicates opposite heading directions (180°).

Using this scalar product *h*_*ij*_(*t*,*τ*), we can compute TDDC, which indicates the alignment between the heading directions of the two agents at various time delays. In order to compensate for high-frequency noise and measurement errors, it is defined as a mean of the heading difference *h*_*ij*_(*t*,*τ*) between *i* and *j* during a symmetric time-window ±*μ t*:

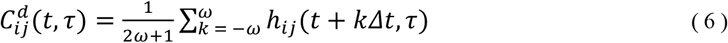

where *t* is the sampling interval, which was set to 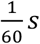 *s* in this study, and *μ* > 0 is a suitably selected constant determined by the reaction time. In this study, *μ* was set to 40 to satisfy *μ t* = 0.66s < 1 s so the heading difference was averaged over an interval less than the follower’s visual-locomotor response time to a leader’s change in heading, which has been estimated as approximately 1 s in walking humans (*35*). Averaging *h*_*ij*_(*t*,*τ*) over a relatively small symmetric time window helps to compensate for measurement noise and tracker error.

To identify the leader-follower relationship, the time delay *τ*_*ij*_(*t*) that maximizes the heading correlation 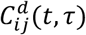 is computed for each pair of agents *i* and *j* at time instant *t*. The sign of the time delay indicates which of the pair is leading the other at time *t:* if *τ*_*ij*_(*t*) has a positive value, it implies that agent *i* was leading agent *j* at the time *t*, and vice versa. Time delay *τ* ranged from -3 s to 3 s. This limit was selected to encompass slow visual-locomotor responses, which can be as much as 2.5s (*35*).

Fig. 2 shows an example heat map characterizing the TDDC values. This example represents the leader-follower interactions between two pedestrians in a 7.83 s segment. The horizontal axis represents time *t*, and the vertical axis represents time delay *τ*. The color temperature of the heat map represents the TDDC value 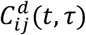, where dark red indicates a TDDC of 1 (i.e., parallel alignment), and dark blue indicates a TDDC of 0 or below (i.e., there is no meaningful correlation between the two headings). The dashed white contour represents a TDDC value of 0.95, and the dashed black line is where there is no time delay (*τ* = 0). Most importantly, the solid white line represents the *τ*_*ij*_(*t*) value that maximizes the heading correlation 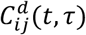 as a function of time. At each time *t* where the solid white line is above the black dashed line (*τ* > 0), it suggests that pedestrian *i* was leading pedestrian *j* at that time. A TDDC value is computed for each possible pair of pedestrians in a sample of crowd data.

### Network reconstruction method

With knowledge of the time delay *τ*_*ij*_(*t*) that maximizes the heading correlation for all pairs of pedestrians, we can infer leader-follower relationships in pedestrian pairs and reconstruct networks illustrating the interactions. In each network snapshot, nodes represent individual pedestrians. Edge direction represents which pedestrian leads the other, and edge weights represent the duration of influence. Specifically, an edge directed from pedestrian *i* to pedestrian *j* indicates that *i* is leading *j*. The edge weight is defined as the percentage of time (number of frames) in which *τ*_*ij*_(*t*) > 0 during the time interval represented by the network snapshot, indicating that *i* is leading *j* for that proportion of the snapshot. A higher weight reflects a longer period of influence.

The reconstructed network represents pedestrian interactions during a given time interval, and after testing various intervals (Fig. S5) we set this interval as 0.5 s. By reconstructing a network snapshot for each consecutive time interval, we can analyze the network dynamics, or how leadership relations evolve over time as individuals change their positions and heading directions.

After all edges in the network with weights > 0 are reconstructed, false-positive connections between nodes are evaluated based on two pruning criteria. (1) By the visibility criterion, a network edge is pruned if the corresponding neighbor is occluded (Fig. S2). Pedestrians are represented as ellipses with a major axis of 0.45 m and a minor axis of 0.244 m (*43*), with the latter aligned with the heading direction. A 180° field of view is centered on the heading direction. The visibility of pedestrian *i* to pedestrian *j* ranges from 0 (fully occluded or out of view) to 1 (fully visible), and it is computed for all ordered pairs at every time point. An edge directed from *i* to *j* is pruned if the mean visibility of *i* to *j* is < 0.15, that is, if the mean proportion of the full ellipse that is visible is less than 0.15 during the time interval of a snapshot (see Fig. S2). This value was determined experimentally by showing that *i* does not influence *j* if *i* is below this cutoff (*44*). (2) By the response time criterion, an edge is pruned if the corresponding time delay is less than 300 ms, because the correlation is likely to be coincidental given actual visual-locomotor response times (*35*).

The reconstructed networks, which we refer to as *visual influence networks*, are embedded in a metric space by plotting the location of each node in the physical position of the corresponding pedestrian at the beginning of the represented time window. The vertical axis represents the group mean heading at the beginning of the 1 s time window. The normalized plot reveals how leadership changes with a pedestrian’s spatial position within the crowd.

### Leadership analysis

We estimate leadership of each pedestrian using four measures that each characterize different features of the visual influence networks. After the network reconstruction, these measures are computed for every pedestrian in each network snapshot.

#### Direct influence

*Direct influence (DI*) assesses the local influence of an individual by summing the outgoing influence on adjacent (directly connected) nodes, minus the incoming influence from adjacent nodes (see Equation 1). The DI of pedestrian *i* is defined as the sum of outgoing edge weights from node *i* (i.e., the sum of pedestrian *i*’s influence on others) minus the sum of ingoing edge weights to node *i* (i.e. the sum of others’ influence on pedestrian *i*). The resulting value is normalized by dividing by the maximum DI value from all pedestrians in the group. The normalization ensures that the DI value for each pedestrian *i* ranges up to 1.

#### Branching influence

Branching *influence (BI*) estimates the magnitude of a given pedestrian’s global influence on the rest of the crowd along all outgoing simple paths (with no repeated nodes), taking both direct and indirect connections into account (see Equation 2). The total influence exerted by pedestrian *i* (a source node) on pedestrian *j* (a destination node) is calculated by taking the product of weights along each possible simple path connecting *i* to *j* (*45*), and then summing the products for all the paths to all pedestrians *j*. This value is subsequently normalized by the maximum BI value observed among all pedestrians in the group. The result captures the overall influence of pedestrian *i* on other pedestrians in the crowd.

#### Direct binary influence

Direct *binary influence (DBI*) of pedestrian *i* assesses a structural property of the network at the local level, rather than estimating the magnitude of influence (see Equation 3). It is similar to DI, but it does not take edge weights into account, only the number of edges (degree). The DBI is defined as the number of outgoing edges from node *i* (outdegree) minus the number of ingoing edges to node *i* (indegree). This value is normalized by dividing by the maximum DBI value for all pedestrians in the group, ensuring that the DBI value for each pedestrian has a maximum of 1.

#### Branching binary influence

*Branching binary influence (BBI*) assesses a structural property of the network at a global level, rather than estimating the magnitude of influence (see Equation 4). It is similar to BI, but it does not take edge weights into account, only the number of outgoing edges along simple paths that originate in a given node. To compute the BBI of a given pedestrian *i*, first we find the total number of simple paths directed from *i* to another pedestrian *j*. This number is summed for all other pedestrians in the network, and then normalized by the maximum BBI value in the group. This creates the BBI range of [0, 1].

### Spatial analysis of leadership

To produce spatial heat maps representing leadership patterns across all networks, we need to normalize positions for the varying spatial sizes and shapes of different groups in different trials (e.g. Fig. S6). To achieve this, we estimate the standard deviations of position along the Front-Back axis and Left-Right axis across trials (the vertical axis represents the mean heading of pedestrian in each network).

We first estimate the spatial distribution of all the datapoints, where each datapoint represents a pedestrian in any network snapshot, and identify spatial outliers. Datapoints appearing in grid cells (0.5 m x 0.5 m) that contain fewer than 5 datapoints are defined as outliers. Removing them at this stage of analysis yields spatial heat maps that do not overrepresent cells with very few datapoints.

A 2D probability density function (PDF) of occupancy (0.5 m x 0.5 m cells) is then computed on the remaining data for each group (Fig. S7), and fitted with a 2D Gaussian function (least squares method) to produce standard deviations (SDs) along the two axes (see the Supplementary Text for the results per group size). The positions of all pedestrians are converted from meters to z-scores and plotted in a coordinate system with units of SD (Fig. S7A). This allows us to aggregate networks across various group sizes and crowd shapes in a standardized coordinate system.

To estimate the spatial distribution of each influence measure in a crowd, we plotted the mean influence values across all networks in a standardized heat map (Fig. 5). The center of mass of the crowd is placed at the origin (0,0), and each cell in the map represents the area of 0.25 SD x 0.25 SD.

## Acknowledgments

Thanks to the Sayles Swarm crew for the data collection; to Trenton D. Wirth and Gregory C. Dachner for providing the analysis scripts; to Maria Lombardi for providing the scripts to calculate the TDDC; and to Matthew Harrison for the conception of branching influence and his thoughtful feedback on the network analysis.

## Data and materials availability

All data are available in the main text or the supplementary materials. Code can be accessed via: https://doi.org/10.5281/zenodo.13765410, and pedestrian position data can be accessed via: https://doi.org/10.26300/xm9v-tg76.

