## Supplementary Information for "Visual Influence Networks in Walking Crowds"

### Supplementary information for
### “Visual Influence Networks in Walking Crowds”

Kei Yoshida,* Mario di Bernardo, William H. Warren

**This PDF file includes:**

Supplementary Text
Figs. S1 to S8

**Supplementary Text**

Normalization of pedestrian positions in spatial heat maps

To produce spatial heat maps in a normalized space, after eliminating spatial outliers, we computed meters per one standard deviation (SD) for the two axes:

- $N = 10$ group: 0.89 m/SD horizontally, 1.30 m/SD vertically
- $N = 16$ group: 1.19 m/SD horizontally, 1.67 m/SD vertically
- $N = 20$ group: 1.39 m/SD horizontally, 1.93 m/SD vertically

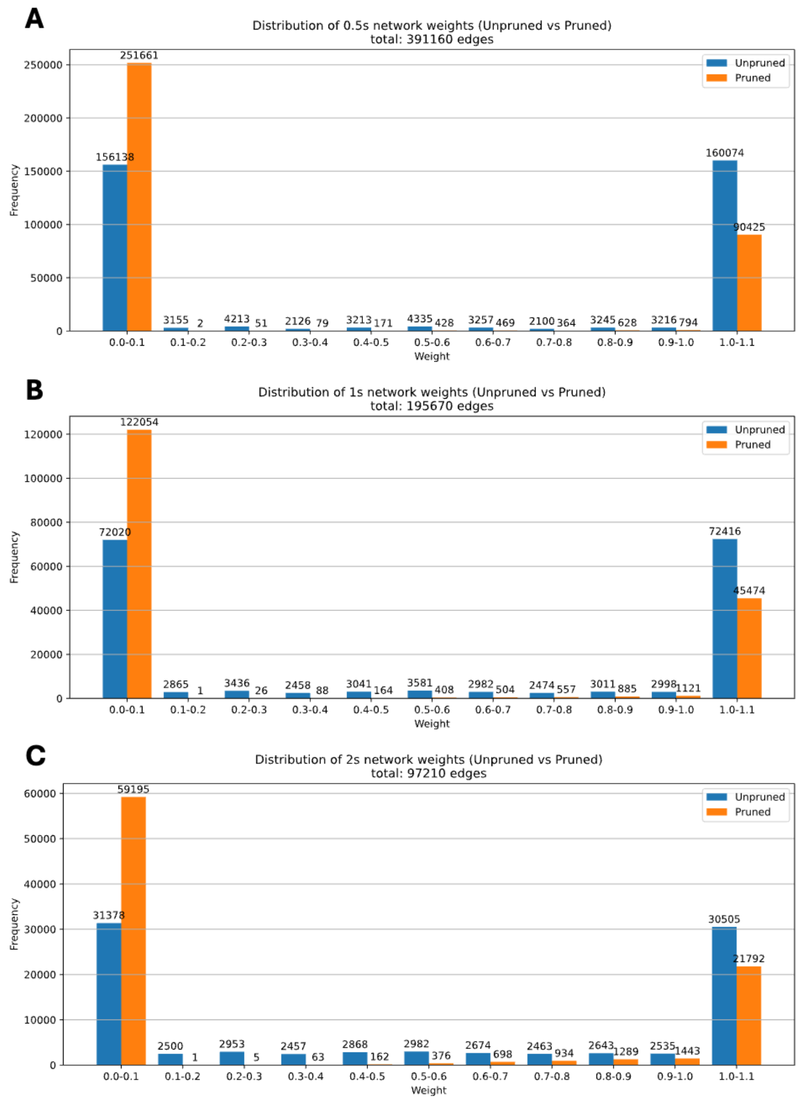


**Fig. S1.** Distribution of network weights for different time window sizes of **(A)** 0.5 s, **(B)** 1 s, and **(C)** 2 s. The blue and red bars respectively represent the weight distribution in the networks before (blue) and after (orange) pruning was applied. The vast majority of weights are close to 0 or 1.

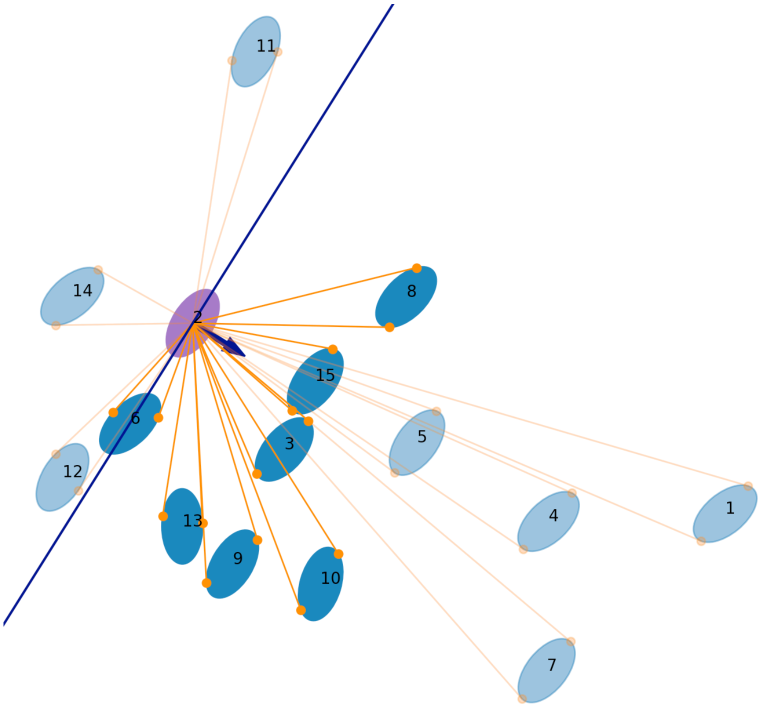


**Fig. S2.** Pruning method of network connections based on visibility. At this instant (N = 16 group), Pedestrian #2 (focal pedestrian shown in purple) is moving in the direction of the thick arrow toward bottom right, with a 180˚ field of view delimited by the purple line. Pedestrians visible to Pedestrian #2 are shown in dark blue. They may be fully or partially visible (≥0.15). Other pedestrians who are not visible to Pedestrian #2 are shown in light blue, due to full or partial occlusion (<0.15), or being outside the field of view.

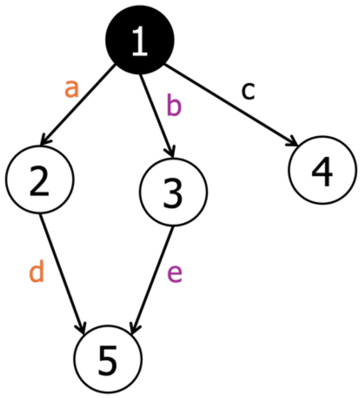


$$BI_{1\_2} = a$$
$$BI_{1\_3} = b$$
$$BI_{1\_4} = c$$
$$BI_{1\_5} = a \times d + b \times e$$

$$BI_1 = BI_{1\_2} + BI_{1\_3} + BI_{1\_4} + BI_{1\_5}$$
$$= a + b + c + (a \times d + b \times e)$$

**Fig. S3.** An illustration of how Branching Influence (BI) is computed for a given pedestrian $i = 1$. The influence of Pedestrian #1 on all pedestrians j ($BI_{1\_j}$) is computed by taking the product of edge weights along each simple path *i,j,* and then summing over all paths. The value is normalized by the maximum BI value observed in this network.

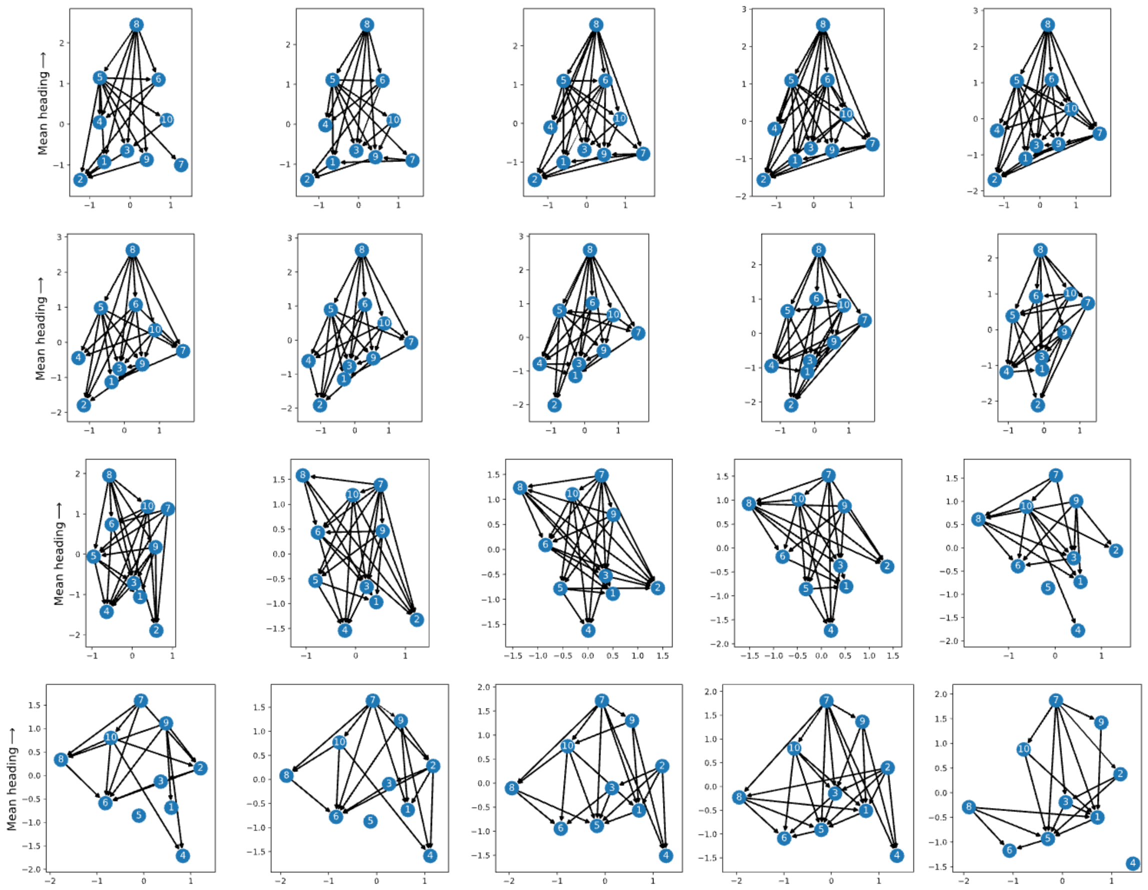

**Fig. S4.** Pruned networks representing the same 10 s time window in Fig. 1 and 4, arranged chronologically from left to right, starting at the top left corner and progressing to the bottom right corner. Participant #8 was at the front in the first 5 s, and Participant #7 take over the front. In Fig. 4, the strongest influence change from #8 to #7. Fig. 3 is 5 networks (network 10-14) taken from this figure where the leadership change occurs.

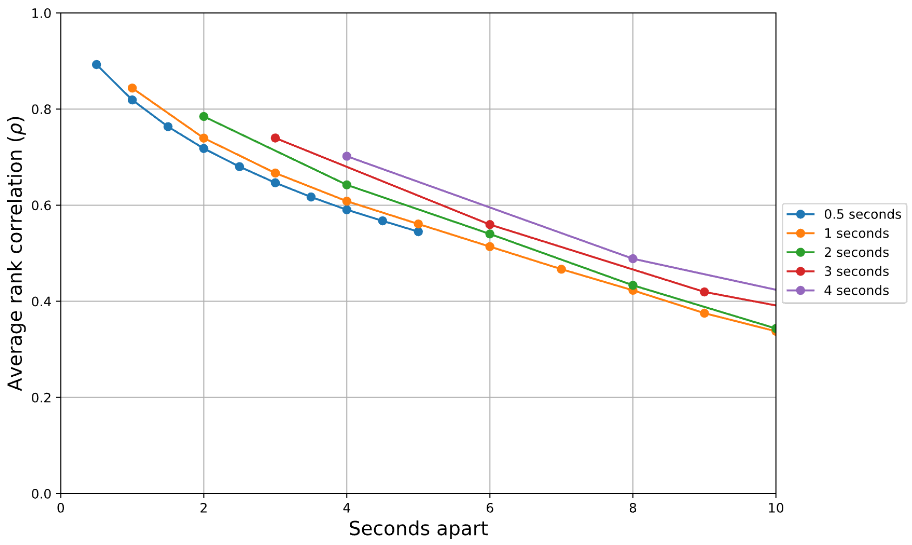


**Fig. S5.** A relationship between average DI rank correlation and network separation in terms of time (in seconds). Curves represent the temporal window of the network reconstruction.
To characterize leadership dynamics, we estimated the rate of change in the DI ranking by comparing networks computed in different time windows (0.5, 1, 2, 3, and 4 s) and separated by different time intervals (number of seconds between networks). The similarities between the rankings in two networks from the same segment of data were determined using the Spearman's rank correlation. The average rank correlation coefficients ($\rho$) are plotted as a function of time between the two compared networks appears in this figure. The correlation between consecutive networks was the highest ($\rho = .90$) for 0.5 s networks, and it remained higher than the correlations for 1-, 2-, 3-, and 4-second networks at all separations. Moreover, the correlation decreased monotonically with temporal separation. This indicates that network topology was highly stable over a period of 0.5 s, but changes markedly over an interval of 2 s, suggesting that a 0.5 s snapshot is appropriate for capturing the rapidly evolving leadership dynamics in the present ‘swarm’ data.

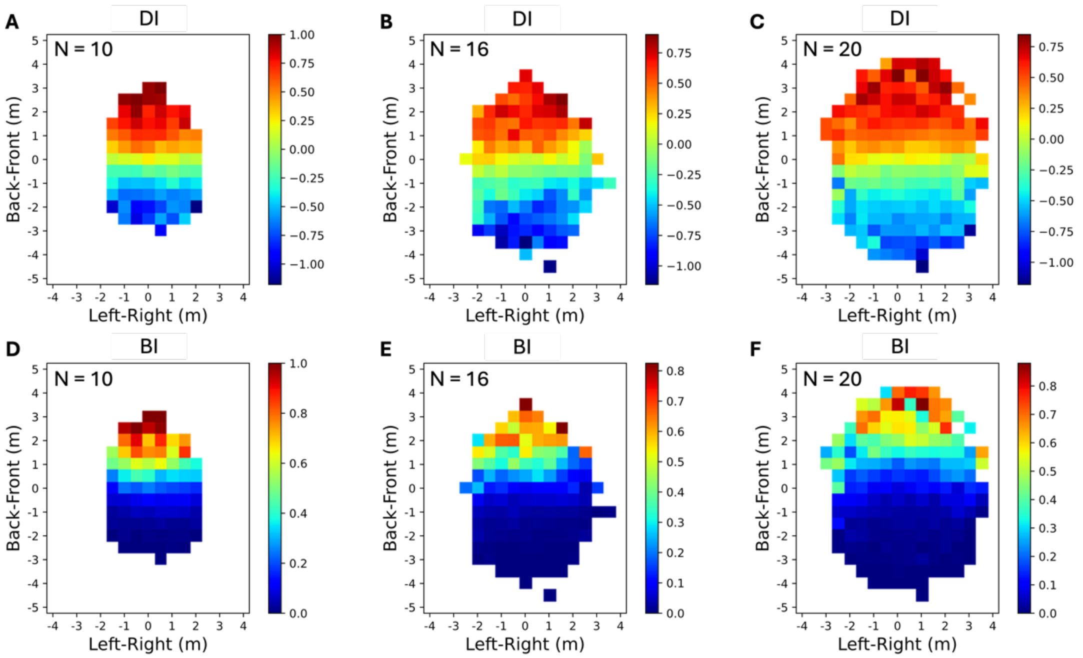


**Fig. S6.** Spatial heat maps of influence without spatial normalization. The axis unit is in meters on both axes. Top row: Net Influence after pruning for **(A)** N = 10, **(B)** N = 16, and **(C)** N = 20 groups. Bottom row: Cumulative Influence for **(D)** N = 10, **(E)** N = 16, and **(F)** N = 20 groups.

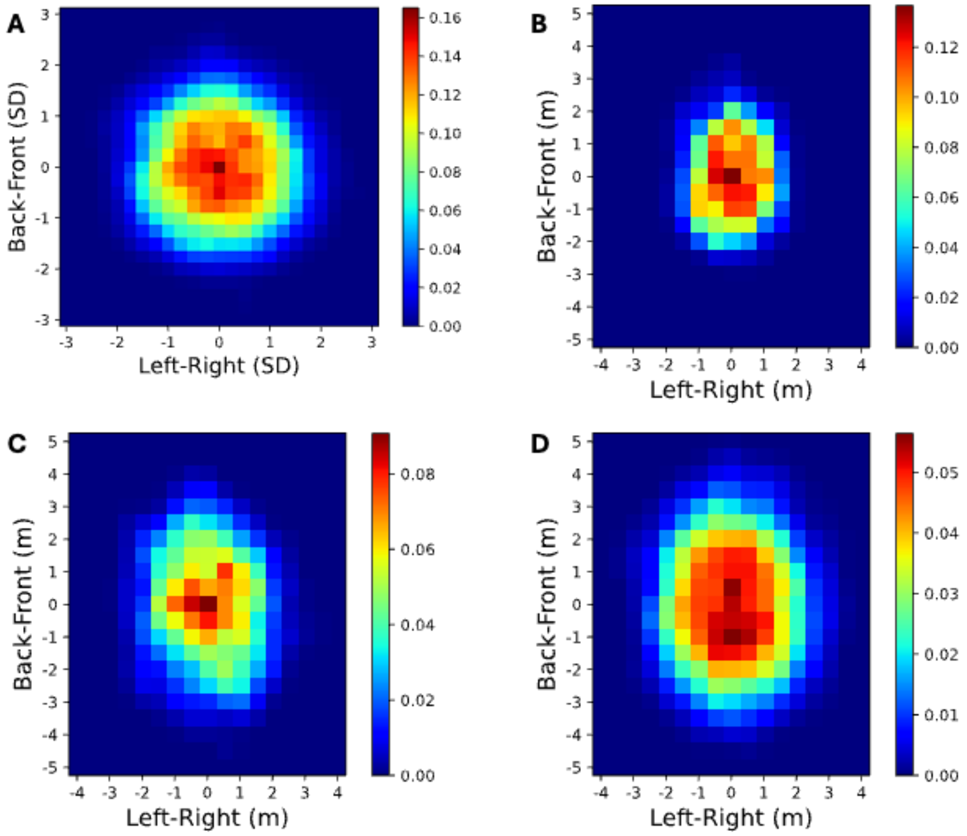


**Fig. S7.** Probability density of occupancy for **(A)** all 3 groups combined, **(B)** N=10 group, **(C)** N=16 group, and **(D)** N=20 group. The unit is in standard deviation (SD) for panel A, and in meters for panels B, C, and D.

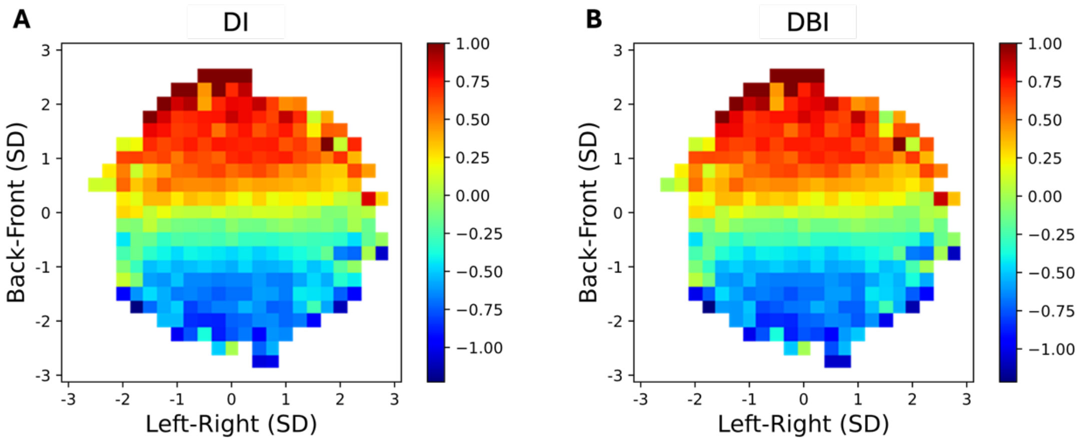


**Fig. S8.** Spatial heat maps of influence before pruning was applied for (a) DI and (b) DBI. The DI and DBI heat maps, both representing local influence, are quite similar after pruning (Fig. 5) to that before pruning (Fig. S5). This similarity indicates that network reconstruction based on TDDC captured the general influence pattern before considering the visual connectivity among pedestrians. Nevertheless, after pruning spurious ingoing and outgoing edges, the heat map has a smoother gradient pattern, with a more homogeneous region of high influence in the front of the crowd, shading into a more localized area of low influence at the rear.
